# Transplanting increases the leaf production rate in rice, consequently modifying plant development and growth

**DOI:** 10.1101/2021.03.08.434354

**Authors:** Benoit Clerget, Estela Pasuquin, Rene Carandang, Abigail J. Domingo, Heathel L. Layaoen, Crisanta Bueno

## Abstract

In Asia, direct sowing and water savings are major trends in previously transplanted and flooded irrigated rice systems because of the higher cost of wages and increasing water shortage. Previous experiments showed that the leaf appearance rate varies between aerobic and flooded cropping systems. This study aimed to further understand how the planting method affects the development rate, flowering time, and yield of lowland irrigated rice crops. A two-year experiment was undertaken at the International Rice Research Institute, Philippines, using three contrasting rice varieties and three planting methods (transplanted, wet direct-seeded, and dry direct-seeded; at a density of 25 plants m^-2^) in a field submerged in 3–5 cm water from two weeks after the transplanting date. The effect of the planting method was similar in the three varieties, mostly without interaction between the two factors. In 2013, the leaf number of seedlings grown in seedling trays was two leaves behind that of direct-seeded plants at the time of transplanting. However, the young transplanted plants recovered quickly; produced new leaves at a faster rate (with a shorter phyllochron); reached panicle initiation, flag-leaf emergence and flowering time 1 week later; and developed more leaves compared to the direct-seeded plants. In 2014, growing in the nursery induced no delay in leaf appearance due to temperatures lower than those in 2013; therefore, the planting method did not affect the leaf appearance rate. Thus, plant development was primarily delayed by the density stress in the seedling trays under warm temperatures; however, the transplanted rice plants had lower plastochron duration than the direct-seeded plants, which made up for the initial delay in leaf appearance. In both years and at similar plant density, the transplanted plants produced more tillers bearing larger upper leaves that led to a higher leaf area index; however, grain yields were similar for the transplanted and direct-seeded crops.

**Highlights:**

- In seedling trays, leaf appearance stopped at the appearance of the fourth leaf.
- After transplanting, leaf appearance resumed at a faster rate than in direct-seeded plants.
- Transplanted plants had delayed panicle initiation and flowering time, more tillers, and more and larger leaves per tiller, but similar grain yield compared to direct-seeded plants at similar planting density.

## 1. Introduction

Historically, rice, the most important crop in Asia, was first dry sown (Pandey and Velasco, 2002). Then, with the intensification of cropping, transplanting became the prevailing planting method. By the 1950s, transplanting was the dominant planting method. However, in recent decades, direct seeding has again become a prominent alternative in Asian rainfed and irrigated lowlands in response to limited water and labour resources. In 2000, the direct-seeded area covered 21% of the total rice-growing area in Asia.

Only experiments in which early-transplanted crops were compared with direct-seeded crops and those in which both crops were managed under standing water were selected in the following literature review. In irrigated tropical lowlands, the change to direct seeding often caused some yield loss (Huang et al., 2011; Kar et al., 2018; Pandey and Velasco, 2002; Rana et al., 2014; Saharawat et al., 2010; Tabbal et al., 2002; Torres et al., 1993), was neutral for yield in some experiments (Kar et al., 2018; Sudhir-Yadav et al., 2011), and led to higher grain yield only in rare cases (Tabbal et al., 2002). In irrigated sub-tropical and temperate lowlands, grain yields of direct-seeded crops were similar to those for transplanted crops in Japan and China (Chen et al., 2014; Liu et al., 2015; Okami et al., 2013); however, yields were higher in Korea (Heu and Kim, 1997). Farmers switched to direct sowing only when it better addressed their socio-economic constraints (Pandey and Velasco, 2002).

In tropical lowlands, crop duration is generally shorter in direct-seeded crops owing to the transplanting shock that transplanted plants experience, which temporarily pauses plant development (Rana et al., 2014; Saharawat et al., 2010; Tabbal et al., 2002; Torres et al., 1993). However, Sudhir-Yadav et al. (2011) did not report any duration difference between the planting methods in plots irrigated with standing water. Conversely, the crop duration was not affected by the planting method in sub-tropical and temperate irrigated lowlands (Chen et al., 2014; Liu et al., 2015; Okami et al., 2013) or was even longer in case of direct-seeded crops in Korea (Heu and Kim, 1997).

Tillering was shown to start and cease later in transplanted crops, compared to the case for direct-seeded crops, leading to higher tiller density and leaf area index (LAI) at maximum tillering (Huang et al., 2011; Sudhir-Yadav et al., 2011; Torres et al., 1993). At harvest, efficient tiller density was either similar or higher in direct-seeded crops than in transplanted crops.

At panicle initiation and flowering time, nitrogen uptake was lower in direct-seeded crops than in transplanted crops (Chen et al., 2014; Liu et al., 2015; Torres et al., 1993). Leaf nitrogen content, soluble protein content, net photosynthetic rate, soluble sugar content, and glutamine synthetase activity were significantly lower in direct-seeded plants than in transplanted plants (Huang et al., 2011).

A study comparing the development and growth of plants in direct-seeded aerobic rice crops and puddled-transplanted flooded crops showed that the leaf appearance rate was significantly slower in aerobic crops than in flooded crops (Clerget et al., 2014). When the confounding factors were studied separately, the aerobic water management and high plant density only marginally decreased the leaf appearance rate, compared to flooded water and low plant density (Clerget et al., 2016; Clerget and Bueno, 2013). The present study aimed to quantify the effect of the planting method on the leaf appearance rate and development and growth in flooded rice crops. Detailed monitoring of the development and growth of three contrasting rice genotypes in dry and wet direct-seeded and puddled transplanted conditions was undertaken.

## 2. Materials and Methods

### 2.1 Site description

Two dry season experiments were conducted from December 2012 to April 2013 and from January to May 2014 at the research farm at the International Rice Research Institute (IRRI), Los Baños, Philippines (14°11’ N, 121°15’ E, 21 m elevation). The topsoil contained 17.1 g organic carbon kg^-1^, 17.4 g total nitrogen (N) kg^-1^, with a pH (CaCl2) of 6.5, cation exchange capacity of 25.4 meq/100 g, and particle size distribution of 58% clay, 33% silt, and 9% sand.

### 2.2 Preliminary checking of the plastochron-phyllochron synchrony

Nemoto et al. (1995) reported that the plastochron and phyllochron always appeared equally in rice, and after the seedling stage, the number of hidden developing leaves inside the sheaths equalled four. However, at the end of the vegetative phase, the plastochron is slightly shorter than the phyllochron and this initial scheme is lost (Nemoto and Yamazaki, 1993). This well-established relationship was checked in 2011 in a tropical indica rice variety bred by the IRRI (NSIC Rc222). In the present study, the plants were sampled weekly from a flooded plot following the methodology described by Clerget et al. (2014). Pre-germinated seeds were sown in seedling trays on 5 January 2011 and were manually transplanted to a puddled-field 13 days later at a distance of 20 × 20 cm between the plants. Starting from 21 days after sowing (DAS) until 70 DAS, five consecutively tagged plants were collected. The number of already appeared leaves was recorded on the main tiller of each plant, which was dissected to record the number of leaves already initiated by the stem apex.

### 2.3 Experimental design and crop management

During each of the two dry seasons, the experiment was performed in a split-plot, randomised complete block design with three replications. The crop establishment method was assigned as the main plot, and the variety as the subplot. Three crop establishment methods were tested: dry direct-seeded (DDS), wet direct-seeded (WDS), and transplanted (TPT). The three varieties used were IR72, a popular irrigated and rainfed variety released in 1988; NSIC Rc222, a high-yielding variety released in 2010 with good adaptation to irrigated and rainfed environments; and SACG7, a Green Super Rice variety from the Chinese Academy of Agricultural Sciences.

The field was ploughed uniformly with the following differences in preparation: the DDS plots were dry-prepared (rotavator), whereas the WDS and TPT plots were water-saturated and puddled. For each experiment, dry direct seeding in the DDS plots and 2-day seed soaking for the WDS and TPT plots commenced on the same dates (5 December 2012 and 8 January 2014). Four dry seeds per hill were manually sown in the DDS plots and four pregerminated seeds per hill were manually sown two days later in the WDS plots. For the TPT plots, pre-germinated seeds were sown in seedling trays (3560 seeds m^-2^, in 1.6 × 1.7 × 3.0 cm individual cells) and were manually transplanted to the TPT plots 16 days later at one seedling per hill. The DDS and WDS plots were thinned to one plant per hill. The distance between each plant was set at 20 × 20 cm, with one plant per hill in all crop methods. The soil was kept saturated in all plots until two weeks post-transplanting and then flooded with 3–5 cm of daily refilled water until two weeks before harvest.

Phosphorus (40 kg P ha^-1^ as single superphosphate), potassium (40 kg K ha^-1^ as muriate of potash), and zinc (5 kg Zn ha^-1^ as zinc sulphate heptahydrate) were applied and incorporated in all plots 1 day before sowing in the 2013 experiment and one day before transplanting in the 2014 experiment. A total of 160 kg N ha-^1^ in the form of urea was applied in four equal splits during the crop cycle. Weeds were controlled manually as required, and pests were controlled using chemicals. A 0.7-m high plastic rat barrier was installed around all plots to prevent damage caused by rats.

### 2.4 Measurements

Weather data were recorded from a station installed at one field extremity. The incident global solar radiation (GS1 dome solarimeter, Delta-TDevices, Cambridge, UK), air temperature at 2 m above ground level (HMP45C, Vaisala, Helsinki, Finland), and soil temperature at a depth of 2 cm (T thermo-couples, PyroControle, Vaulx en Velin, France) were measured at 1-min intervals, averaged on an hourly basis, and stored in dataloggers (CR1000, Campbell Scientific, Logan, UT, USA). Soil temperature was recorded in the nursery, using three thermo-couples in three randomly chosen cells in three seedling trays and with the same dataloggers.

Before the transplanting date, leaf appearance was recorded every 2–3 days in 2013 for five plants from each of the 18 directly sown plots and for five plants from 3 seedling trays of each variety in the nursery. The same was undertaken daily in 2014 in six directly sown plots and in one tray per variety in the nursery. Immediately after transplanting, 16 sampling rows were randomly identified in each of the 27 plots. Reserved 5-m^2^ harvest areas were also marked on the same day. In each sampling row, seven consecutive plants were marked with a coloured ring and each blade of the main tiller of these plants was tagged based on its order of appearance. The first incomplete leaf, or prophyll, was counted as leaf 1 (Yoshida, 1981).

Starting at the transplanting day, five consecutive plants from the pre-marked sampling rows were uprooted weekly from each plot. The plant samples were carefully washed to remove the soil and transported to the laboratory for plant processing. The numbers of tillers and fully expanded leaves, appeared leaves, and senesced leaves, and plant height (height from soil surface to the highest visible collar) of the main tiller were recorded for each plant. A new tiller was counted when its first leaf emerged from its enclosing sheath.

The area, length, and width of each fully expanded, individually tagged green leaf from the main tiller were measured (LI-3100C Area Meter, Li-Cor, Lincoln, NE, USA). A leaf was considered fully expanded when its collar became visible above the enclosing sheath of the previous leaf. Similarly ranked leaves were later grouped and stored in paper bags. Dead leaf blades, leaf sheaths, culms, and panicles (including juveniles when present) from the five main tillers from the plot samplings were grouped into organ-specific bags. The remainder of the tillers, green leaf blades, dead leaf blades, leaf sheaths, culms, and panicles (including juveniles when present) were separated into specific bags after the leaf area of the green blades was recorded. All samples were dried for 72 h in an oven at 70°C prior to weighing. Total plant leaf area was calculated as the sum of the individual leaves from the main tiller and the green blades of the other tillers. The LAI was calculated as the leaf area of the sampled plants divided by the corresponding ground sampling area. The shoot dry matter of the plants was calculated as the sum of the dry matter of the leaf blades, leaf sheaths, culm, and panicles.

In the 2013 experiment, five additional plants per plot were sampled at mid-tillering, panicle initiation, flowering, and harvest, and were dried at 60°C for 72 h. Aliquot samples from the stems, leaf blades, and panicles were extracted and milled. The N content was then analysed using the Kjeldahl method at the IRRI-Analytical Service Laboratory.

Flowering time was determined in each plot when an average of 50% of the spikelets per panicle of the main tillers of 50% of the observed plants had extruded their anthers. Crop maturity was established when 95% of the spikelets of the entire plot had turned yellow.

At maturity, the grains were separated from the rachis and the filled and unfilled grains were separated at a flow rate of 4 m^3^s^-1^ using a Seedburo blower (KL-1205, Seedburo, Chicago, IL, USA). The total weight of the filled and unfilled grains and the weight of 1000 filled and unfilled grains were measured. The moisture content of the filled grains was measured using a digital moisture tester (DMC 700, Seedburo). The number of filled and unfilled grains was estimated via the ratio of the total weight to the weight of 1000 grains, whereas the fertility rate was computed as the ratio of the filled grains to the total number of grains. The harvest index was computed as the filled grain dry matter divided by the shoot dry matter. The grain yield at 14% moisture content was estimated both from the 5-plants processed at harvest time and from the 5-m^2^ sampling area reserved in the centre of each plot.

### 2.5 Data analysis

Thermal time (TT) was calculated from the day of emergence of the seedlings as the cumulative sum of the mean daily temperature of the apical meristem minus the base temperature of 11°C. Until the onset of stem elongation, when stems longer than 5 cm started to constitute a separate sampled organ, the meristems were close to the soil surface and their temperature was estimated to be equal to the soil temperature at a depth of 2 cm. Once the stems were longer than 5 cm, the meristem temperature was estimated using the air temperature.

The leaf appearance rate was assumed to follow trilinear kinetics, or a broken-line model with three segments, in which the first four leaves appeared faster than the succeeding leaves (Clerget and Bueno 2013). Thus, the observed leaf number (LN) of the main stem was regressed against the elapsed TT from plant emergence using the following equation:

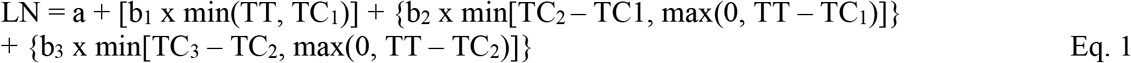

where a is the number of leaves at emergence; b_1_, b_2_, and b_3_ are the initial, secondary, and tertiary development rates, respectively; TC_3_ is the TT when the last leaf appeared; and TC1 and TC_2_ are the TTs when the development rate changed. The parameters and their confidence intervals were iteratively estimated using the NLIN procedure (SAS, 2012). Phyllochron1, Phyllochron2, and Phyllochron3 were calculated as the inverse of b_1_, b_2_, and b_3_, respectively.

In rice, many Japanese scientists have shown that leaf initiation and appearance curves are parallel, stably distant by four leaves (Nemoto et al. 1995), and slightly diverging later (Nemoto and Yamazaki, 1993). This relationship was checked in 2011 at IRRI on the NSIC Rc222 variety (Fig. 1). The number of developing leaves inside the sheaths was equal to 4 until the initiation of leaf 12 and thereafter progressively increased to 5 until the initiation of the last leaf. The TT at panicle initiation (TTPI) in the main stem was assumed to be synchronous with the appearance of the (N – 5)^th^ leaf and was estimated from the reverted trilinear model, with N being the total number of leaves produced by the stem as follows:

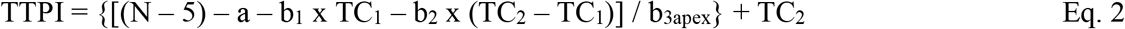

where b_3apex_ = b_3_ x 5/4 is the rate of leaf initiation that is faster than the rate of leaf appearance during the last phase.

**Fig. 1:**
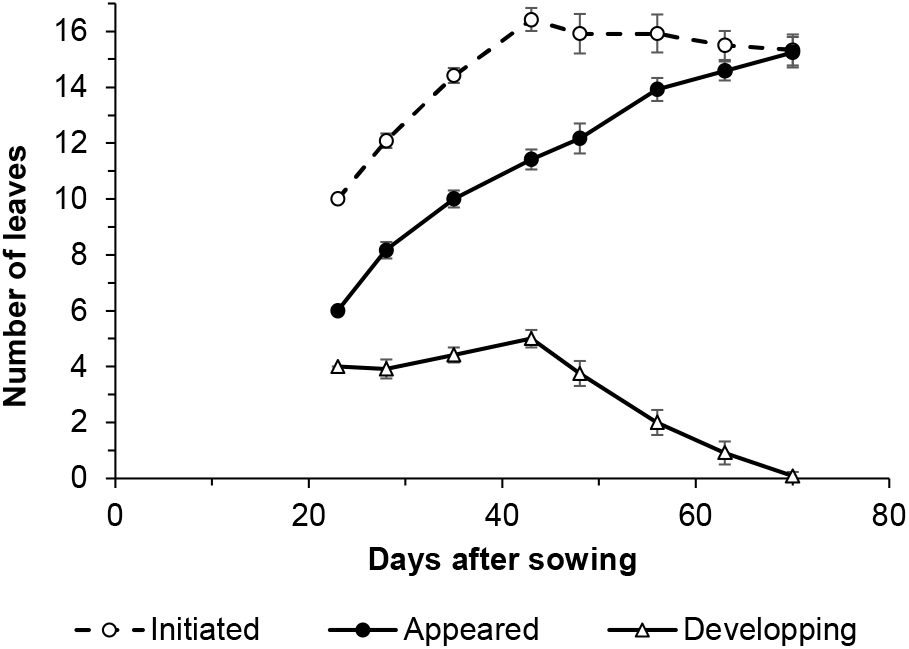
Observed mean numbers of initiated and appeared leaves, and by difference the number of developing leaves on the main stems of rice plants from the NSIC Rc222 variety. Seedlings were transplanted 15 days after sowing and the crop grew in flooded conditions in 2011 at the IRRI’s farm (Clerget et al., 2014). Error bars indicate the 95% confidence interval of each mean number.

The relationship between the number of tillers per plant and the number of leaves on the main stem was modelled as a numerical series. Two series were tested: nbtil_n_ = nbtil_n-1_ + nbtil_n-2_ (Fibonacci series) and nbtil_n_ = nbtil_n-1_ + nbtil_n-3_, where nbtiln is the number of tillers at the appearance of the n^th^ leaf on the main stem. The number of leaves on the main stem at the time of the onset of tillering (n_onset_) was iteratively estimated using the Solver function in Microsoft Excel, and the series that best fit the observations was selected (lower root mean square deviation).

Analysis of variance was performed using the general linear model procedure (GLM, SAS, 2012) with supplementary Tukey’s pairwise mean comparison.

Plant nitrogen content was estimated for the four sampling dates (mid-tillering, panicle initiation, flowering, and harvest) and was compared to the critical N% dilution curve drawn as a function of shoot biomass (BM):

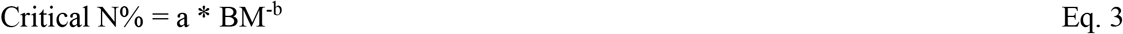

where parameter values calculated for rice by Sheehy et al. (1998) from flooded rice plots fertilised with 165 kg ha^-1^ N and grown in the Philippines (a = 3.74 and b = 0.44) were used.

## 3. Results

### 3.1 Climate

The weather was exceptionally dry during the first four months of 2014 because of the onset of the intense 2014–2016 El Niño event (Yin et al., 2018). Additionally, January 2014 was cooler than usual, with a monthly mean temperature of 24.3°C, which was 1°C below the 1979–2013 long-term average. Consequently, the daily average temperatures were noticeably higher during the first month of cultivation in the 2013 experiment than in the 2014 experiment (Fig. 2A). Except during the first 2 weeks of cultivation, the daily solar radiation was higher in 2014 than in 2013 (Fig. 2B). The cumulative rainfall was 42 mm in 2014, compared to 382 mm in 2013 (Fig. 2C).

**Fig. 2:**
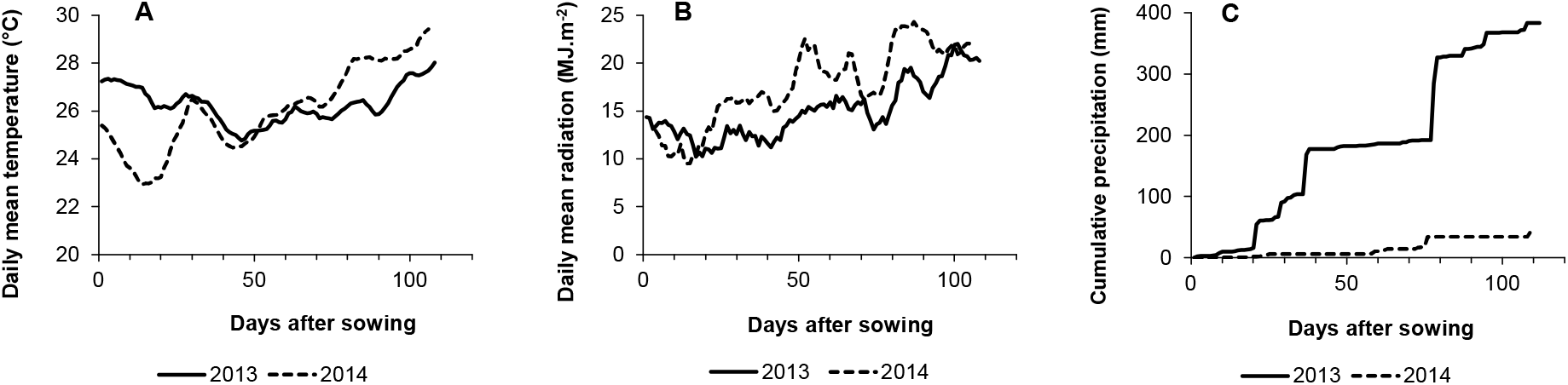
Ten days moving average of daily mean temperature (A) and daily global sun radiation (B), and cumulative precipitations (C) during the 2013 and 2014 dry seasons.

### 3.2 Leaf appearance rate and plant phenology

In 2013, large differences in the leaf appearance kinetics were observed between the direct-seeded plants and transplanted plants, whereas WSD and DDS plants had similar kinetics (Fig. 3A, Supplementary Fig. 1A and C, Table 1, and Supplementary Table 1).

**Fig. 3:**
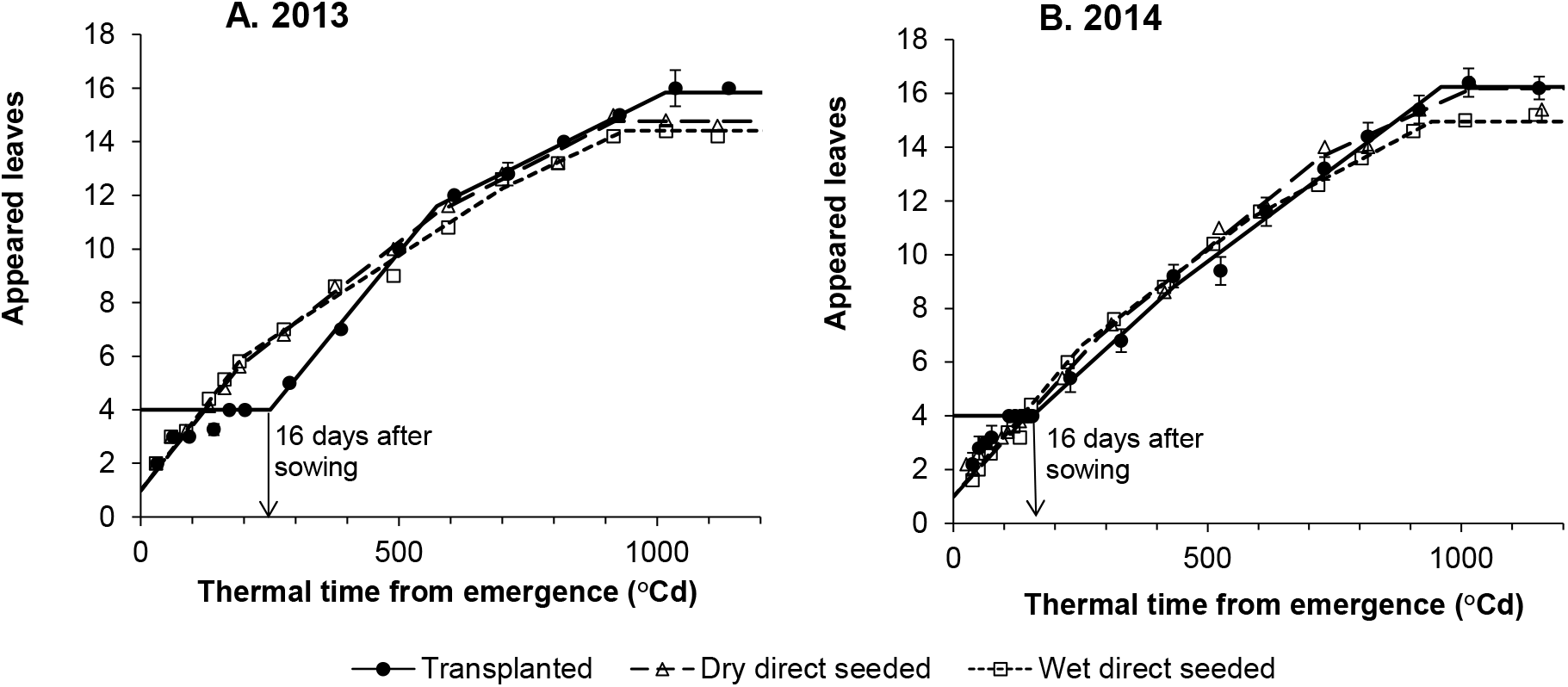
Mean observed number (symbols) and predicted number (lines) of appeared leaves on the main stem of plants from the variety NSIC Rc222 per planting method in 2013 (A) and 2014 (B) dry seasons. Arrows show the thermal time at transplanting date. Error bars indicate the 95% confidence interval of each mean number for the transplanted treatment.

**Table 1:**
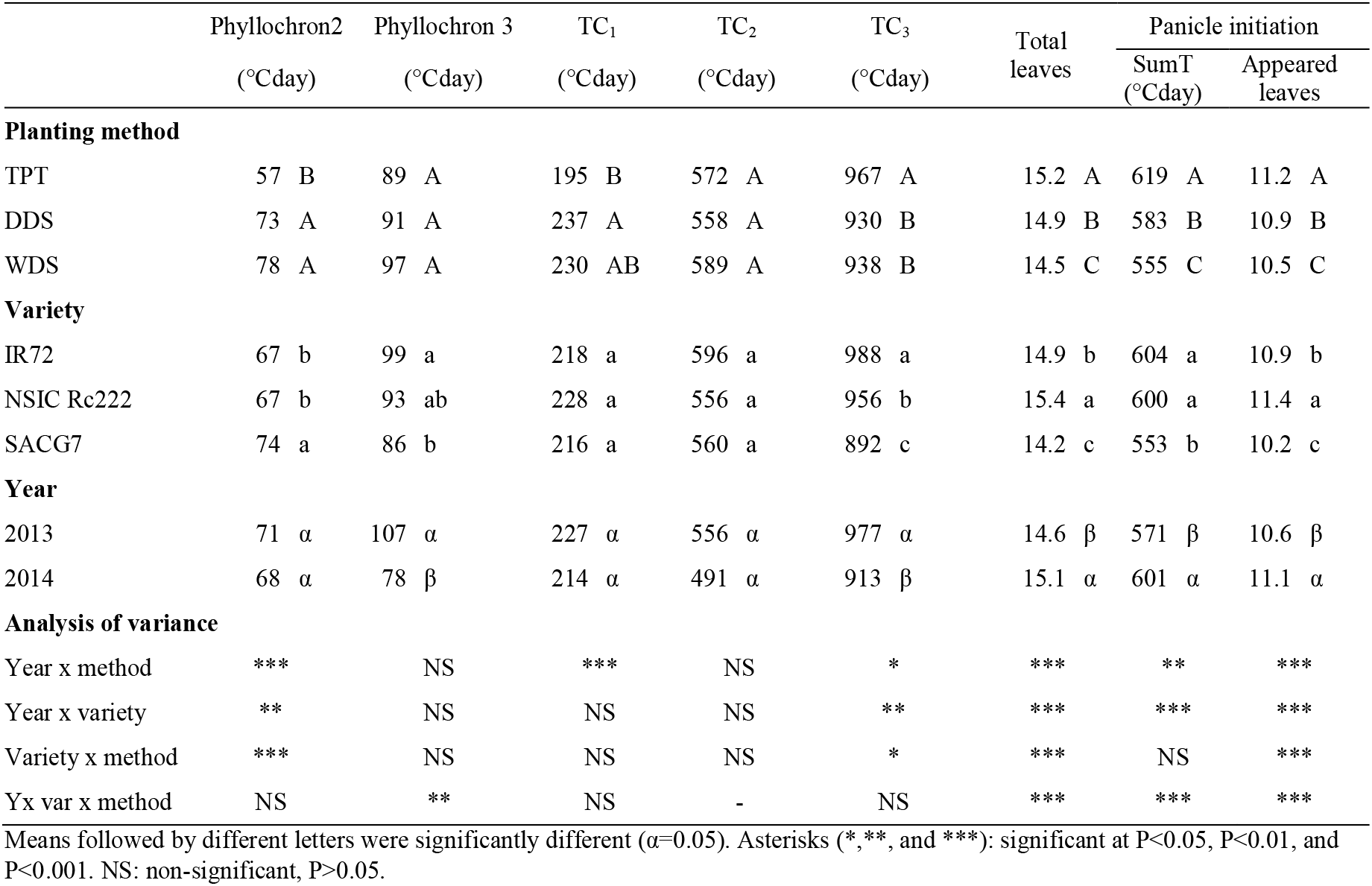
Means per factor of the broken-linear model parameters and of the estimated occurrence of panicle initiation in thermal time and in number of appeared leaves on the main stem, and probabilities of the ANOVA’s F-test for the interactions between factors.

|  | Phyllochron2<br>(°Cday) | Phyllochron 3<br>(°Cday) | TC<sub>1</sub><br>(°Cday) | TC<sub>2</sub><br>(°Cday) | TC<sub>3</sub><br>(°Cday) | Total<br>leaves | Panicle initiation |  |
| --- | --- | --- | --- | --- | --- | --- | --- | --- |
|  |  |  |  |  |  |  | SumT<br>(°Cday) | Appeared<br>leaves |
| **Planting method** |  |  |  |  |  |  |  |  |
| TPT | 57 B | 89 A | 195 B | 572 A | 967 A | 15.2 A | 619 A | 11.2 A |
| DDS | 73 A | 91 A | 237 A | 558 A | 930 B | 14.9 B | 583 B | 10.9 B |
| WDS | 78 A | 97 A | 230 AB | 589 A | 938 B | 14.5 C | 555 C | 10.5 C |
| **Variety** |  |  |  |  |  |  |  |  |
| IR72 | 67 b | 99 a | 218 a | 596 a | 988 a | 14.9 b | 604 a | 10.9 b |
| NSIC Rc222 | 67 b | 93 ab | 228 a | 556 a | 956 b | 15.4 a | 600 a | 11.4 a |
| SACG7 | 74 a | 86 b | 216 a | 560 a | 892 c | 14.2 c | 553 b | 10.2 c |
| **Year** |  |  |  |  |  |  |  |  |
| 2013 | 71 $\alpha$ | 107 $\alpha$ | 227 $\alpha$ | 556 $\alpha$ | 977 $\alpha$ | 14.6 $\beta$ | 571 $\beta$ | 10.6 $\beta$ |
| 2014 | 68 $\alpha$ | 78 $\beta$ | 214 $\alpha$ | 491 $\alpha$ | 913 $\beta$ | 15.1 $\alpha$ | 601 $\alpha$ | 11.1 $\alpha$ |
| **Analysis of variance** |  |  |  |  |  |  |  |  |
| Year x method | *** | NS | *** | NS | * | *** | ** | *** |
| Year x variety | ** | NS | NS | NS | ** | *** | *** | *** |
| Variety x method | *** | NS | NS | NS | * | *** | NS | *** |
| Yx var x method | NS | ** | NS | - | NS | *** | *** | *** |

Means followed by different letters were significantly different ( $\alpha=0.05$ ). Asterisks (\*, \*\*, and \*\*\*): significant at $P<0.05$ , $P<0.01$ , and $P<0.001$ . NS: non-significant, $P>0.05$ .

The plants sown in the nursery showed a slow appearance of the fourth leaf that was the latest to appear before transplanting (Fig. 4A). In contrast, dry direct-seeded plants already developed 5.8 leaves at this time. However, a few days after transplanting, the rhythmical appearance of new leaves of the transplanted plants resumed at a rate that significantly surpassed that of the direct-seeded plants (Phyllochron2 was 36 to 34°Cd shorter) and had produced even more leaves by the time the direct-seeded plants produced their 12th leaf. The leaf appearance rates then slowed down and showed homogenous values among the three planting methods until the appearance of the flag leaf (Phyllochron3). The flag leaf appeared 60 to 69°Cd (TC_3_) or four days later in the transplanted plants after the appearance of 0.8 to 1.0 more leaves than in the direct-seeded plants.

**Fig. 4:**
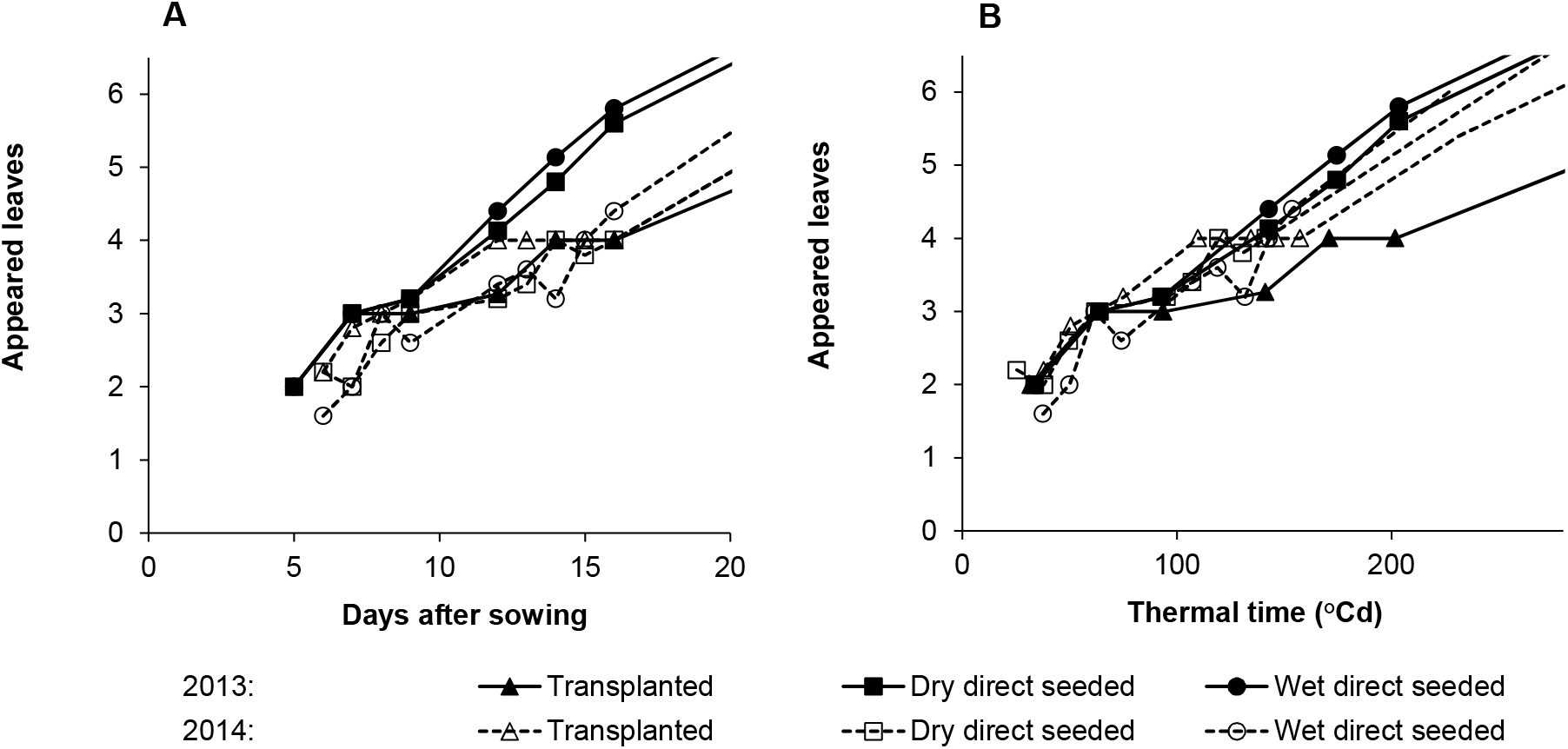
Detail of the number of appeared leaves on the main stem of NSIC Rc222 until the transplanting details (16 days after sowing) against A) days after sowing and B) thermal time since sowing, per year and per planting method.

Similarly, in 2014, plants in the nursery did not develop further after the appearance of the fourth leaf for 5 days (Fig. 3B, Fig. 4A, Supplementary Fig. 1B and D, and Supplementary Table 2). Transplanting was undertaken 16 DAS in both years, but at different TTs; therefore, the 2014 experiment was performed 45°Cd earlier due to the cooler mean temperatures when the plants were in the nursery. This lower temperature also delayed seedling emergence and the appearance of leaves in the direct-seeded plants (Fig. 4B). Consequently, transplanted and direct-seeded seedlings had a similar number of leaves (4) at the time of transplanting in contrast with the situation in 2013, when a difference of 1.5 leaves was observed, and Phyllochron2 was only 7 to 9°Cd shorter in transplanted plants.

Three of the model parameters (Phyllochron2, TC_1_, and TC_2_) had stable means between the years (Table 1). Conversely, TT, the number of appeared leaves at panicle initiation, and total number of leaves on the stem were significantly lower in 2013 than in 2014. However, the duration to flag leaf appearance (TC_3_) was significantly longer in 2013 than in 2014 in relation to the significantly longer Phyllochron3.

The IR72 and NSIC Rc222 varieties had close estimates of the model parameters. SACG7 showed significantly earlier panicle initiation and flag leaf appearance (TC_3_), with fewer total leaves produced by the main stem.

### 3.3 Tillering

Significantly more tillers per plant were produced until maximum tillering in 2014 than in 2013; however, there was no difference between the years at harvest time (Fig. 5A, Supplementary Fig. 2A and G, and Table 2). Plants of the IR72 variety produced significantly more tillers than the NSIC Rc222 plants and even more tillers than the SACG7 plants. However, at harvest time, only SACG7 had a significantly lower number of tillers among the three varieties. The number of tillers per plant increased exponentially with TT in both years; however, the tiller outgrowth stopped earlier in 2013 than in 2014. The onset of tillering was delayed slightly with the transplanted plants and tillering ceased 1 week later, producing significantly more tillers than the direct-seeded plants; however, the difference was not apparent at harvest time.

**Fig. 5:**
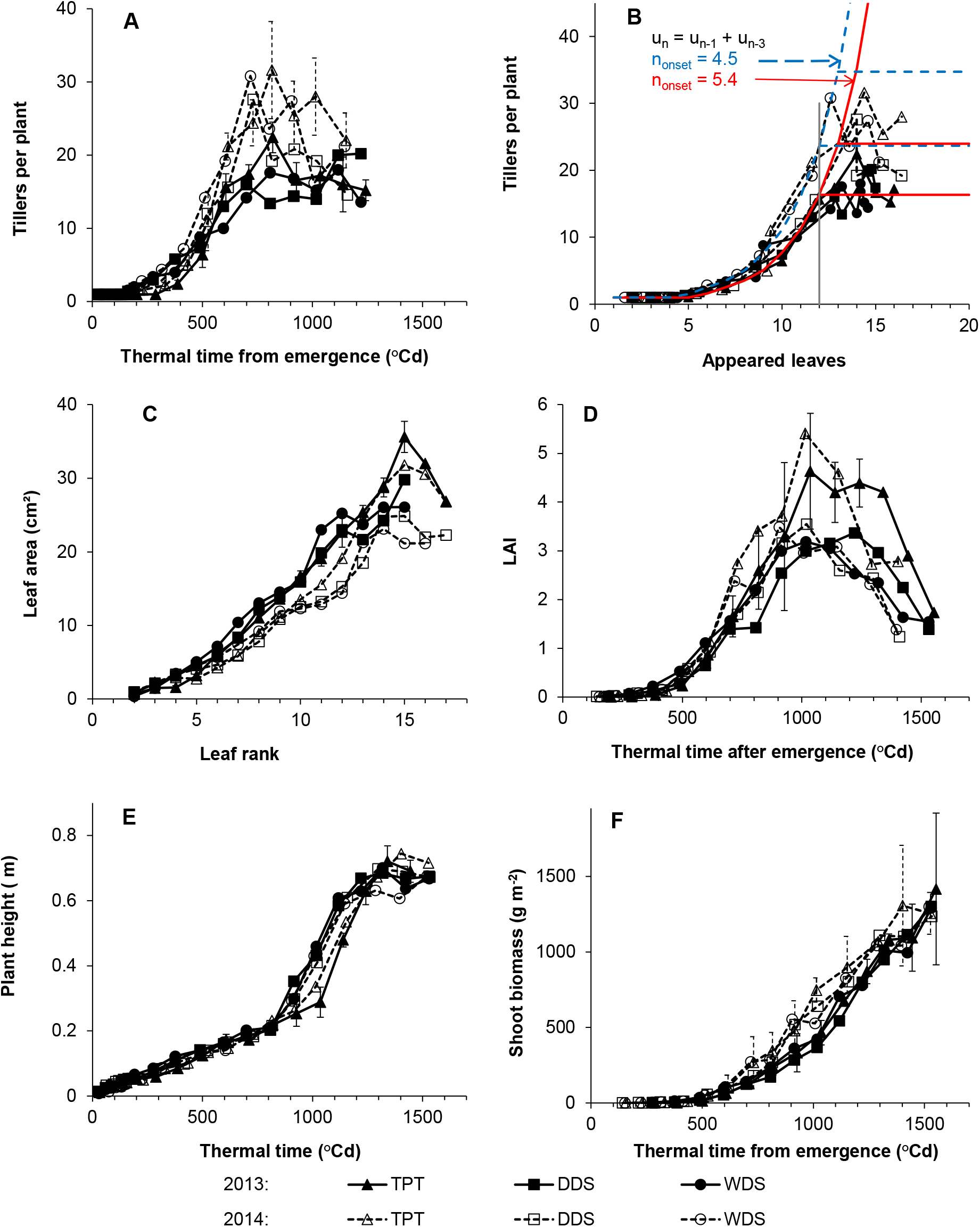
Mean tillering with time (A) and with leaf appearance (B), areas of the leaves of the main stem (C), LAI dynamics (D), plant height (E), and shoot biomass accumulation (F) for the NSIC Rc222 planted in three methods in 2013 and 2014. Error bars indicate the 95% confidence interval of each value for the transplanted treatments.

**Table 2:** Means of tillering, leaf area of leaves until rank 12, final plant height, and LAI traits per factor and probabilities of the ANOVA’s F-test for the interactions.

|  | Maximum<br>tiller number | Tillers at<br>harvest | Mean area of<br>leaves <=12 | Final plant<br>height (m) | Maximum<br>LAI | SumT at<br>max LAI | LAI at<br>harvest |
| --- | --- | --- | --- | --- | --- | --- | --- |
| **Planting method** |  |  |  |  |  |  |  |
| TPT | 23.4 A | 12.3 A | 12.6 C | 0.70 A | 4.6 A | 1110 A | 2.4 A |
| DDS | 20.4 B | 11.3 A | 14.0 B | 0.67 B | 3.5 B | 1117 A | 1.5 B |
| WDS | 20.5 B | 11.8 A | 15.3 A | 0.66 C | 3.6 B | 1049 A | 1.5 B |
| **Variety** |  |  |  |  |  |  |  |
| IR72 | 29.8 a | 15.4 a | 11.2 b | 0.60 c | 4.3 a | 1077 a | 2.0 a |
| NSIC Rc222 | 25.1 b | 14.2 a | 9.9 c | 0.68 b | 4.1 a | 1077 a | 1.7 a |
| SACG7 | 9.4 c | 5.7 b | 20.1 a | 0.77 a | 3.3 b | 1124 a | 1.6 a |
| **Year** |  |  |  |  |  |  |  |
| 2013 | 17.4 $\beta$ | 11.6 $\alpha$ | 16.8 $\alpha$ | 0.68 $\alpha$ | 4.0 $\alpha$ | 1155 $\alpha$ | 1.9 $\alpha$ |
| 2014 | 25.5 $\alpha$ | 12.0 $\alpha$ | 11.7 $\beta$ | 0.67 $\beta$ | 3.8 $\alpha$ | 1030 $\beta$ | 1.6 $\alpha$ |
| **Analysis of Variance** |  |  |  |  |  |  |  |
| Year x method | ** | *** | *** | NS | * | NS | NS |
| Year x variety | * | NS | *** | NS | ** | NS | * |
| Variety x method | NS | NS | *** | *** | ** | NS | NS |
| Y x var x method | NS | * | *** | ** | NS | NS | NS |

Means followed by different letters were significantly different ( $\alpha=0.05$ ). Asterisks (\*, \*\*, and \*\*\*): significant at $P<0.05$ , $P<0.01$ , and $P<0.001$ . NS: non-significant, $P>0.05$ .

The numerical series nbtil_n_ = nbtil_n-1_ + nbtil_n-3_ well fitted the internal relationship between the numbers of tillers and leaves on the main tillers except for the IR72 variety in the transplanted and WDS treatments in 2013 (Fig. 5B and Supplementary Fig. 2B and H). Therefore, the newly appeared tillers only started to outgrow their own tillers after three more leaves had appeared on the main stem. For NSIC Rc222, tillering started earlier in the plant development in the TPT and WDS plots in 2013 (n_onset_ = 4.5) than in 2014 and in DDS in 2013 (n_onset_ = 5.4); however, it ceased soon after the appearance of the 12th leaf of the main stem in all situations. Thus, the longer developmental duration in TPT and WDS in 2013 was responsible for the greater number of tillers than in the other treatments. For SACG7, late n_onset_ (5.9 and 6.2) and early cessation of the tiller outgrowth between the appearance of the 10th and 11th leaves led to smaller numbers of tillers. For IR72, the relationship between the numbers of tillers and appeared leaves followed a Fibonacci series (nbtil_n_ = nbtil_n-1_ + nbtil_n-2_) in the TPT and WDS in 2013 whereas n_onset_ was early (4.3) and tillering ceased between the appearance of the 11th and 12th leaves in all treatments. Consequently, the IR72 plants produced a higher number of tillers than plants of the other varieties.

### 3.4 Growth: Leaf size, LAI, plant height, and biomass accumulation

The mean area of leaves up to rank 12 on the main stem was significantly larger in 2014 than in 2013, significantly larger in the WDS than in the DDS and TPT treatments, and significantly different among the varieties (Fig. 5C, Supplementary Fig. 2C and I, and Table 3). In the transplanted plants, significantly more leaves were produced by the main stem and the upper leaves were larger than in the direct-seeded plants (Table 1). Because of the larger leaf area per stem and the higher number of tillers per plant, maximum LAI values were significantly higher in the transplanted than in the direct-seeded crops in both years (Fig. 5D, Supplementary Fig. 2D and J, and Table 2). The onset of stem elongation of the transplanted plants occurred 1 week later than in the direct-seeded plants in case of all varieties in 2013 and only in NSIC Rc222 in 2014 (Fig. 5E and Supplementary Fig. 2E and H). The year and planting method had significant but small effects on the final plant height, whereas the different varieties had significantly different heights (Table 2).

**Table 3:** Mean yields and yield components per factor and probabilities of the ANOVA’s F-test for the interactions between factors.

|  | Shoot biomass<br>(g m<sup>-2</sup>) | N content 2013<br>(%) | Grain Yield (g m<sup>-2</sup>) |  |  | Harvest index | Panicle density<br>(m<sup>-2</sup>) | Spikelets per panicle | Fertility rate<br>(%) | Grain weight<br>(mg) |
| --- | --- | --- | --- | --- | --- | --- | --- | --- | --- | --- |
|  |  |  | 5-plants samples | 5-m<sup>2</sup> samples | Difference samples |  |  |  |  |  |
| Planting method |  |  |  |  |  |  |  |  |  |  |
| TPT | 1239 A | 1.07 A | 672 A | 670 A | -2 A - | 48 B | 287 A | 132 A | 73 B | 26.9 A |
| DDS | 1172 A | 0.89 B | 666 A | 608 B | -58 A + | 50 AB | 267 A | 135 A | 77 AB | 26.9 A |
| WDS | 1202 A | 0.89 B | 701 A | 624 B | -77 A + | 51 A | 269 A | 133 A | 80 A | 27.1 A |
| Variety |  |  |  |  |  |  |  |  |  |  |
| IR72 | 1318 a | 0.93 a | 771 a | 697 a | -73 b | 52 a | 349 A | 103 c | 82 a | 26.4 b |
| NSIC Rc222 | 1299 ab | 0.98 a | 764 a | 688 a | -78 b | 52 a | 334 a | 118 b | 76 b | 25.9 b |
| SACG7 | 997 b | 0.94 a | 505 b | 520 b | +15 a | 45 b | 140 b | 179 a | 72 b | 28.6 a |
| Year |  |  |  |  |  |  |  |  |  |  |
| 2013 | 1268 α | 0.95 | 713 α | 656 α | -57 α | 49 α | 271 α | 148 α | 74 β | 26.7 β |
| 2014 | 1142 β | - | 646 β | 612 β | -34 α | 50 α | 278 α | 119 β | 79 α | 27.2 α |
| Analysis of variance |  |  |  |  |  |  |  |  |  |  |
| CV% | 14 | 9 | 16 | 5 | 228 | 7 | 15 | 13 | 8 | 2 |
| Year x method | NS | - | NS | NS | NS | NS | ** | ** | NS | ** |
| Year x variety | NS | - | * | *** | NS | NS | NS | NS | NS | *** |
| Variety x method | NS | NS | NS | NS | NS | NS | NS | NS | NS | NS |
| Y x var x method | NS | - | NS | NS | NS | NS | NS | NS | NS | NS |

Means followed by different letters were significantly different ( $\alpha=0.05$ ) and +/- were for significant contrast between groups ( $\alpha=0.05$ ). Asterisks (\*, \*\*, and \*\*\*): significant at $P<0.05$ , $P<0.01$ , and $P<0.001$ . NS: non-significant, $P>0.05$ .

In 2014, the linear phase of shoot biomass accumulation was reached at an earlier TT and stopped 1 week earlier than in 2013 (Fig. 5F and Supplementary Fig. 2F and L). Shoot biomasses at harvest were similar across the planting methods and were significantly lower in 2014 than in 2013 due to the earlier cessation of biomass accumulation (Table 3). The N content of the transplanted plants was at the same level of the dilution curve reported by Sheehy et al. (1998) for a transplanted crop grown in soil fertilised with 165 kg ha^-1^ N at all sampling dates, whereas it was always slightly lower in the direct-seeded plants (Supplementary Fig. 3).

### 3.5 Yield and yield components

In 2013, harvesting was undertaken at plant maturity on 20 March (105 DAS) in 15 plots, on 25 March (110 DAS) in the transplanted NSIC Rc222 plots, and on 27 March (112 DAS) in the IR72 plots. In 2014, all plots were harvested on 28 April (110 DAS). Grain yield estimations from the five plants and 5-m^2^ samples were inconsistent relative to the planting method (Table 3). No significant difference was shown in the five plant samples, whereas the 5-m^2^ samples from the transplanted plots showed significantly higher yields than those from the direct-seeded plots. Yield estimations from the 5-m^2^ samples were less variable (coefficient of variation [CV] = 5%) than those from the five plant samples (CV = 16%). Yield estimations from both sample sizes were consistent in the transplanted plots; however, they were contrastingly lower in the 5-m^2^ samples from the direct-seeded plots. The SACG7 plants showed significantly lesser yields than the two other varieties and showed a significantly lower difference between the estimations from the two sample sizes. The significant interaction between the year and variety was caused by a reversal of the ranking for the yields of NSIC Rc222 and IR72 in both years. The mean grain yield was consistently and significantly lower in 2014 than in 2013.

The fertility rate and harvest index were significantly lower in the transplanted plants, whereas the other yield components were not affected by the planting method (Table 3). A significant difference between the varieties for all yield components was observed. The number of spikelets per panicle, fertility rate, and grain weight varied significantly between the years.

## 4. Discussion

Inter-year variability was large and significant for the majority of the measured traits, yields, and yield components; however, the effects of the two experimental factors (crop planting and variety) were consistent across the years and could therefore be conclusively analysed.

### 4.1 Differential shattering at harvest

The direct-seeded plots and the plants of the SACG7 variety that headed earlier might have matured earlier than the transplanted plots and the two other varieties (Table 3). All plots were harvested on the same date in both years, except for the NSIC Rc222 plants in the transplanted plots and IR72 plants that were harvested one week later in 2013. This difference in the cycle duration would explain the significant negative difference between the grain yield estimations of the 5-m^2^ and five plant samplings in the direct-seeded plots. At harvest, more grains were possibly shattered from the over-matured panicles in the 5-m^2^ samples than from the more carefully managed small five plant samples. The two dwarf IR72 and NSIC Rc222 varieties would shatter more than the taller SACG7 variety in agreement with the known linkage between the sd1 dwarfing gene and sh2 shattering gene (Nakamura et al., 1995; Oba and Kikuchi, 1991). Consequently, grain yield estimations from the five plant samplings that did not respond to the planting method better reflected the potential yields, which would be nearly attainable using a combine-harvester. The yield estimation from the 5-m^2^ plots better reflected the harvesting by the manual labour of farmers.

### 4.2 Plants stressed in the nursery developed faster and for longer after transplanting than the direct-seeded plants

Flag leaf appearance and grain maturity occurred earlier in the direct-seeded plants than in the transplanted plants. This agreed with the general observation in tropical rice crops that has been explained by transplanting shock delaying plant development (Rana et al., 2014; Tabbal et al., 2002; Torres et al., 1993). However, the plants recovered quickly as previously reported by Pasuquin et al. (2008) and Torres et al. (1993). In the present study, new leaves started appearing a few days after transplanting and the difference in the plant development stage was acquired before transplanting. After the first week and the appearance of the third leaf, the development of the seedlings growing in the small cells of the seedling trays slowed down and finally stopped at the appearance of the fourth leaf (Fig. 4A). Seedlings grown under heterotrophic conditions in the dark also stop their development at the fourth leaf stage (Yoshida, 1981). The small space per seedling would prevent the acquisition of autotrophy in the seedling trays. Consequently, in plants transplanted two weeks after sowing from the seedling trays, the developmental delay against the direct-seeded plants was not caused by transplanting shock, but instead, by a nursery effect.

Leaf appearance accelerated stably after transplanting in response to the stress experienced in the nursery. An accumulation of growing leaves inside the sheaths when the plants were in the nursery would have caused the same result. However, previous observations made in Japan where transplanting is a normal practice (Nemoto et al., 1995; Nemoto and Yamazaki, 1993) and confirmed at the farm at IRRI (Fig. 1) showed a stable number of leaves (four) growing inside the sheaths in rice during the beginning of the vegetative phase. Thus, the production rhythm of new leaf primordium by the apical meristem of the stem was modified twice by the environmental conditions: first, in the seedling trays, and second, at the time of transplanting. Variations in the leaf appearance rate caused by environmental factors have been reported previously (Wilhelm and McMaster, 1995). Typically, abiotic stresses such as low phosphorus availability (Pellerin et al., 2000) or constraints such as high plant density (Clerget et al., 2016) caused a decrease in the leaf appearance rate. Conversely, in the present study, the faster leaf appearance rate observed after density stress in the nursery caused active compensation, similar to the increased production of flowers after early insect damage reviewed by Harris (1974) and reported in soybean (Haile et al., 1998), cotton (Castella et al., 2005), and rapeseed (Pinet et al., 2015). In cotton, moderate drought stress could be favourable to both boll yield and fibre quality (Niu et al., 2018).

### 4.3 Transplanting affected the crops until harvest time

The duration until the appearance of the last leaf (TC_3_) was significantly longer in transplanted plants than in the direct-seeded plants (Table 1 and Fig. 3). There was a stable relationship between the numbers of initial and appeared leaves in the rice (Fig. 1) and the appearance rate of the last leaves (Phyllochron3) was similar among the three planting methods; therefore, the later flag leaf appearance rate in the transplanted plants was caused by delayed panicle initiation.

The faster leaf appearance rate in transplanted plants possibly caused this delay in panicle initiation. Indeed, correlations between the phyllochron and duration to panicle initiation under inductive photoperiods have been previously observed in sorghum, although in an opposite way (Clerget et al., 2008; Gutjahr et al., 2013). In a sequential series, because of the longer vegetative phase associated with a faster leaf initiation rate, transplanted plants produced more leaves of larger area for the upper ones, outgrew more tillers, and had a higher maximum LAI than direct-seeded plants (Tables 2 and 3). These results agreed with observations from previous studies (Huang et al., 2011; Sudhir-Yadav et al., 2011; Torres et al., 1993). Additionally, transplanted plants had a significantly higher nitrogen content than direct-seeded plants on all sampling dates (Supplementary Fig. 3), in agreement with previous studies (Chen et al., 2014; Huang et al., 2011; Liu et al., 2015; Torres et al., 1993). However, at harvest time, the number of tillers per plant and the panicle density were only little (and not significantly) larger in transplanted plants compared to the direct-seeded ones, whereas the fertility rate was significantly lower in transplanted plants. This plastic adjustment across the yield components resulted in similar potential grain yields among the three planting methods, in agreement with previous studies (Kumar et al., 2016).

### 4.4 Influence of temperature during the seedling stage on the differences in crop duration among planting methods

In 2013, the duration to flag leaf appearance was 1 week or 100°Cd later in the transplanted plots than in the direct-seeded plots (Fig. 3A). In 2014, the duration to flag leaf appearance of the transplanted plants was only 1.3 days or 18°Cd later than that for WDS plants but 4.2 days or 60°Cd earlier than for DDS plants (Fig. 3B). Flowering and maturity durations were primarily piloted by the duration to flag leaf appearance, with a clear contrast between the transplanted and direct-seeded plants in 2013 compared to those in 2014, similar to the leaf appearance kinetics. The unusually cool temperature during seedling emergence in 2014 reduced and blurred the clear contrast that was observed in 2013.

This observation regarding the effects of temperature in the nursery on crop duration could explain the contrasting reports on the effects of direct seeding on crop duration. The crop duration of direct-seeded crops is generally shorter than that of transplanted crops in tropical areas (Rana et al., 2014; Saharawat et al., 2010; Tabbal et al., 2002; Torres et al., 1993); however, it is similar or even longer in plants grown in sub-tropical and temperate areas (Chen et al., 2014; Heu and Kim, 1997; Liu et al., 2015; Okami et al., 2013). In temperate and sub-tropical areas, rice planting is undertaken as early as possible during spring when the temperature remains cool, especially during the night, leading to similar plant development in the field and nursery. Thus, as observed in the 2014 experiment, crop duration would not be affected by the planting method under the climatic conditions in these areas.

## 5. Conclusions

The transplanting of rice seedlings in puddled paddy fields has been a very efficient method on the way towards rice intensification. The method addressed various issues related with the cropping calendar and the weed control. The present study showed that, in places with warm temperatures during seedling emergence, this planting method relied on the ability of rice plants to adjust their plastochron in response to an abiotic stress and to its release. This active modification of the plastochron allowed the transplanted plants to recover from the density stress in the nursery. At similar plant densities, transplanted plants produced more tillers with larger upper leaves and had a larger maximum LAI; however, there was no yield advantage compared to the case for the direct-seeded plants.

## Supporting information

Supplementary figs and tables to Rice planting methods

## Acknowledgements

We thank Pedro Gapas, Luis Malabayabas, and Victor Lubignan, who were in charge of the crop physiology experimental tasks.

