## Supplementary figs and tables to Rice planting methods for "Transplanting increases the leaf production rate in rice, consequently modifying plant development and growth"

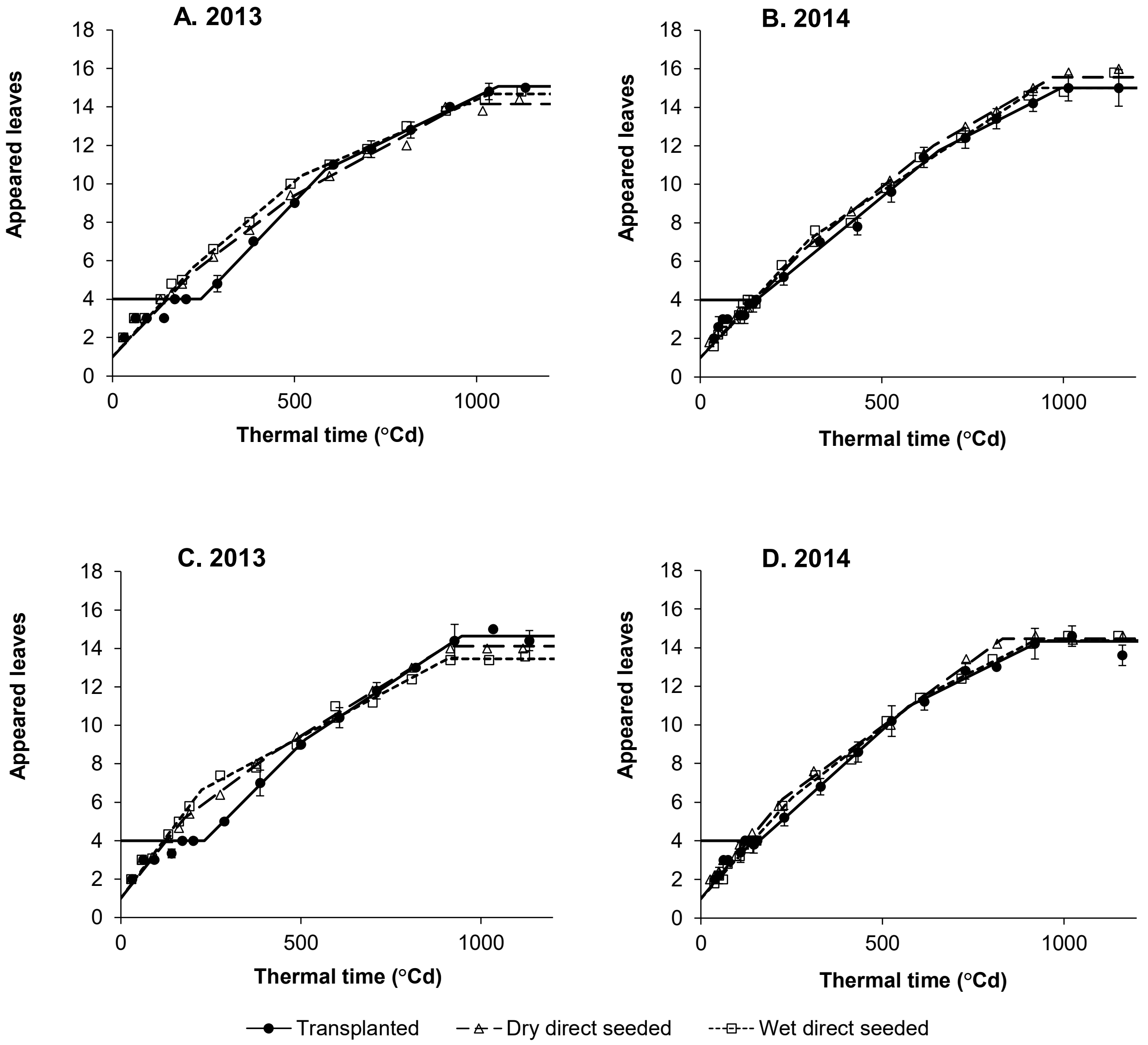


**Supplementary Fig. 1:** Mean observed number (symbols) and predicted number (lines) of appeared leaves on the main stem of plants from the varieties IR72 (A, B) and SACG7 (C, D) per planting method in 2013 (A, C) and 2014 (B, D) dry seasons. Error bars indicate the 95% confidence interval of each mean number for the transplanted treatment.

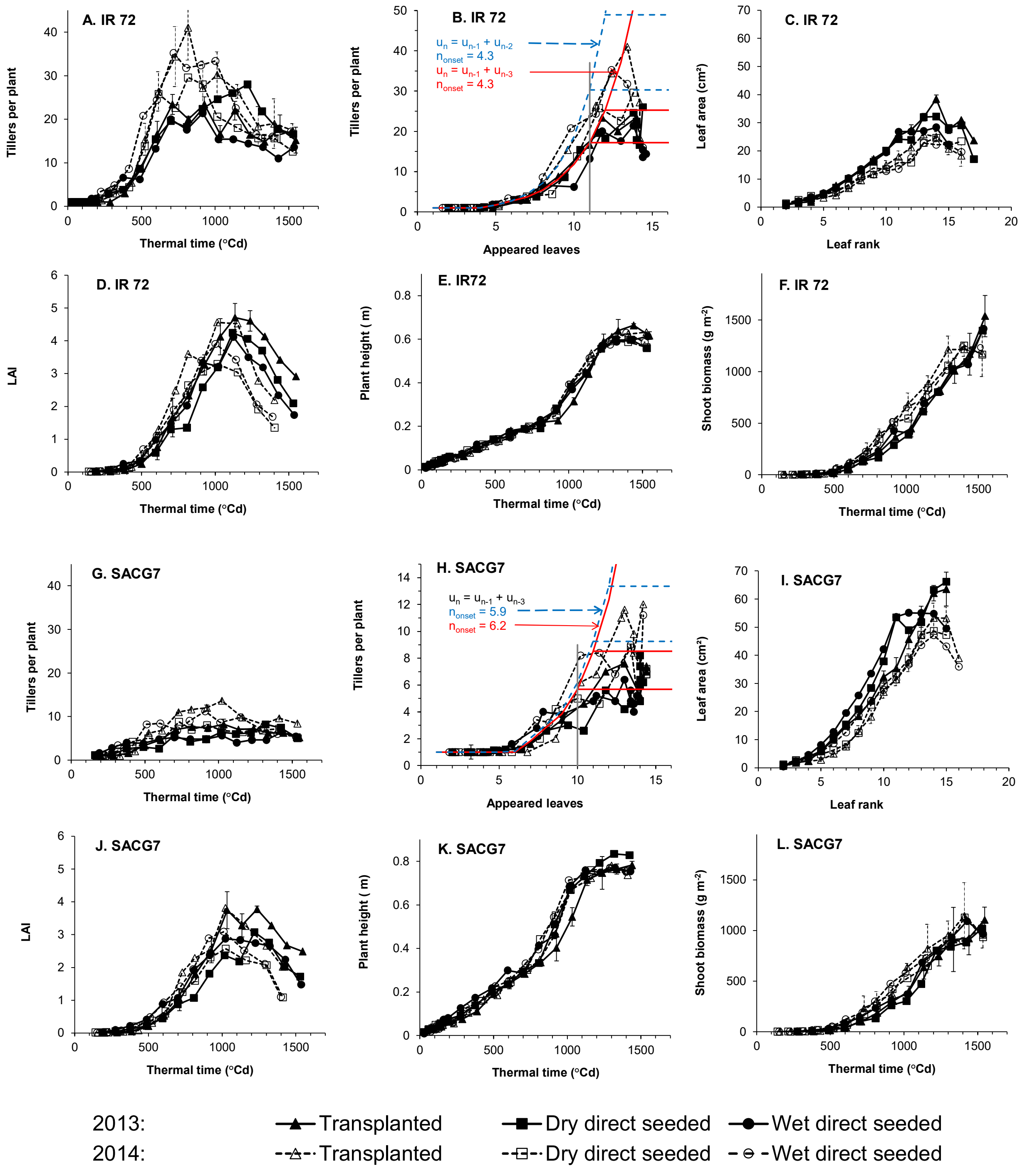


**Supplementary Fig. 2:** Mean tillering with time (A,G) and with leaf appearance on the main stem (B,H), areas of the leaves from the main stem (C,I), and LAI dynamics (D,J), plant height (E,K), and shoot biomass accumulation (F,L) with thermal time for the varieties IR72 and SACG7 planted in three methods in 2013 and 2014. Error bars indicate the 95% confidence interval of each value for the transplanted treatments.

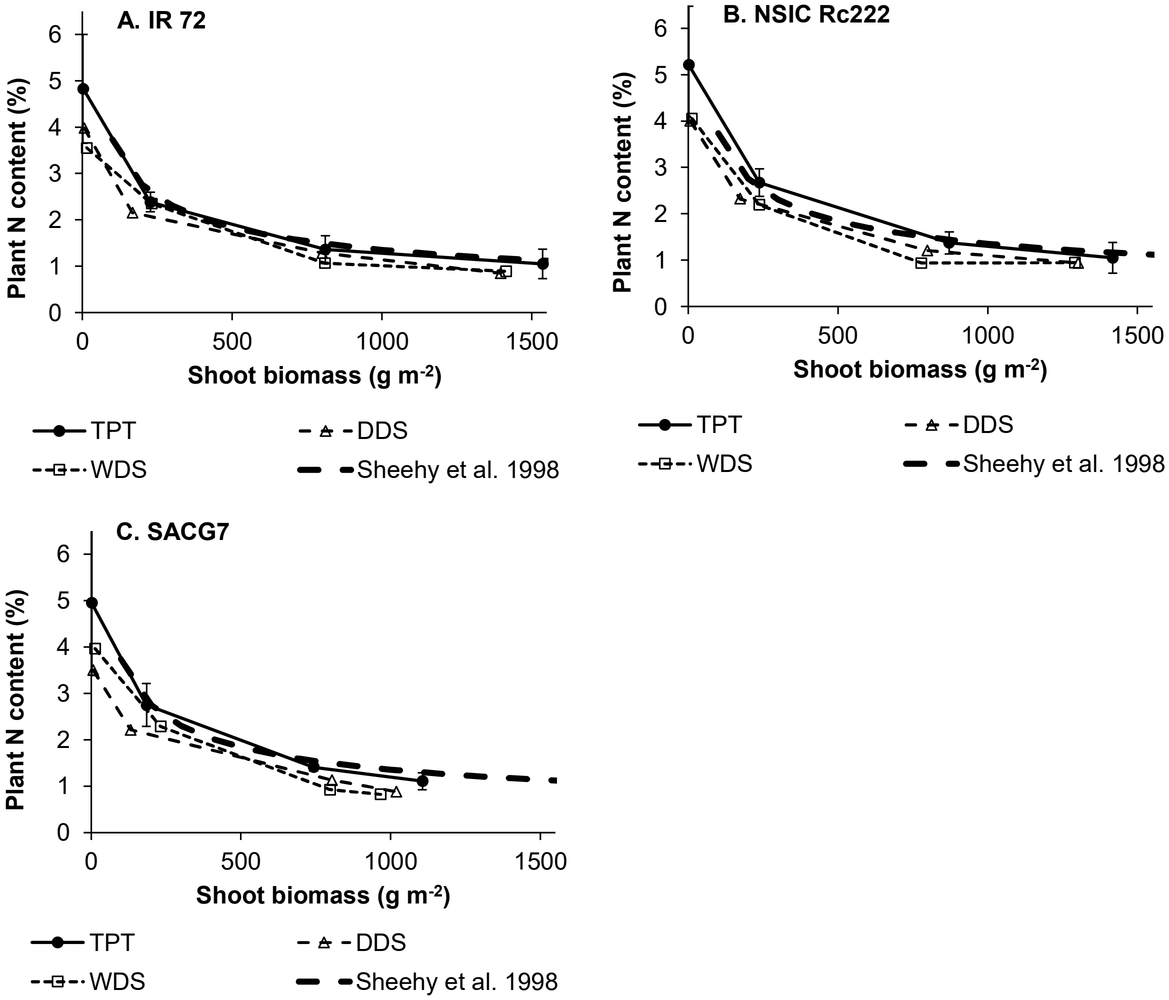


**Supplementary Fig. 3:** The relationship between plant nitrogen content and shoot dry weight compared with the dilution curve established by Sheehy et al. (1998) from flooded crops receiving 165 kg $ha^{-1}$ nitrogen for the three varieties IR72 (A), NSIC Rc222 (B), and SACG7 (C) planted in three methods in 2013. Samples were taken at transplanting, mid-tillering, flowering, and grain maturity. Error bars indicate the 95% confidence interval of each mean value for the transplanted treatments.

**Supplementary Table 1:** 2013 means per factor of the broken-linear model parameters and of the estimated occurrence of panicle initiation in thermal time and in number of appeared leaves on the main stem, and probabilities of the ANOVA's F-test for the interaction between factors.

| | Phyllochron2 (°Cday) | Phyllochron 3 (°Cday) | $TC_1$ (°Cday) | $TC_2$ (°Cday) | $TC_3$ (°Cday) | Total leaves | Panicle initiation | |
|---|---|---|---|---|---|---|---|---|
| | | | | | | | SumT (°Cday) | Appeared leaves |
| **Planting method** | | | | | | | | |
| TPT | 48 C | 105 A | 244 A | 562 A | 1020 A | 15.2 A | 617 A | 11.2 A |
| DDS | 74 B | 106 A | 217 A | 540 A | 952 B | 14.4 B | 561 B | 10.4 B |
| WDS | 82 A | 108 A | 220 A | 540 A | 960 B | 14.2 C | 537 C | 10.2 C |
| **Variety** | | | | | | | | |
| IR72 | 61 b | 118 a | 227 a | 561 a | 1038 a | 14.7 b | 586 a | 10.7 b |
| NSIC Rc222 | 67 ab | 106 ab | 232 a | 584 a | 962 b | 15.0 a | 567 a | 11.0 a |
| SACG7 | 75 a | 96 b | 221 a | 512 a | 933 c | 14.1 c | 559 b | 10.1 c |
| **Analysis of variance** | | | | | | | | |
| Variety x method | * | NS | NS | NS | * | *** | ** | *** |

Means followed by different letters were significantly different (α=0.05). Asterisks (*,**, and ***): significant at P<0.05, P<0.01, and P<0.001. NS: non-significant, P>0.05.

**Supplementary Table 2:** 2014 means per factor of the broken-linear model parameters and of the estimated occurrence of panicle initiation in thermal time and in number of appeared leaves on the main stem, and probabilities of the ANOVA's F-test for the interaction between factors.

| | Phyllochron2 (°Cday) | Phyllochron 3 (°Cday) | $TC_1$ (°Cday) | $TC_2$ (°Cday) | $TC_3$ (°Cday) | Total leaves | Panicle initiation | |
|---|---|---|---|---|---|---|---|---|
| | | | | | | | SumT (°Cday) | Appeared leaves |
| **Planting method** | | | | | | | | |
| TPT | 65 B | 73 B | 146 B | 483 A | 913 A | 15.2 A | 622 A | 11.2 A |
| DDS | 72 AB | 75 B | 256 A | 526 A | 908 A | 15.4 A | 607 A | 11.4 A |
| WDS | 74 A | 86 A | 240 A | 533 A | 916 A | 14.8 B | 572 B | 10.8 B |
| **Variety** | | | | | | | | |
| IR72 | 72 a | 79 a | 208 a | 565 a | 938 a | 15.2 b | 622 a | 11.2 b |
| NSIC Rc222 | 67 a | 80 a | 223 a | 457 a | 950 a | 15.8 a | 632 a | 11.8 a |
| SACG7 | 72 a | 76 a | 211 a | 513 a | 851 b | 14.4 c | 547 b | 10.4 c |
| **Analysis of variance** | | | | | | | | |
| Variety x method | NS | NS | NS | - | NS | *** | *** | *** |

Means followed by different letters were significantly different (α=0.05). Asterisks (*,**, and ***): significant at P<0.05, P<0.01, and P<0.001. NS: non-significant, P>0.05.
